# Pathogen evolution: slow and steady spreads the best

**DOI:** 10.1101/225599

**Authors:** Todd L. Parsons, Amaury Lambert, Troy Day, Sylvain Gandon

**Author notes:** Corresponding author: CEFE UMR 5175, CNRS - Universite de Montpellier-Universite Paul-Valery Montpellier-EPHE, 1919 route de Mende, 34293 Montpellier, France. E-mails: TLP -; AL -; TD -; SG.

## Abstract

The theory of life history evolution provides a powerful framework to understand the evolutionary dynamics of pathogens in both epidemic and endemic situations. This framework, however, relies on the assumption that pathogen populations are very large and that one can neglect the effects of demographic stochasticity. Here we expand the theory of life history evolution to account for the effects of finite population size on the evolution of pathogen virulence. We show that demographic stochasticity introduces additional evolutionary forces that can qualitatively affect the dynamics and the evolutionary outcome. We discuss the importance of the shape of pathogen fitness landscape and host heterogeneity on the balance between mutation, selection and genetic drift. In particular, we discuss scenarios where finite population size can dramatically affect classical predictions of deterministic models. This analysis reconciles Adaptive Dynamics with population genetics in finite populations and thus provides a new theoretical toolbox to study life-history evolution in realistic ecological scenarios.

## 1 Introduction

Why are some pathogens virulent and harm their hosts while others have no effect on host fitness? Our ability to understand and predict the evolutionary dynamics of pathogen virulence has considerable implications for public-health management (Dieckmann et al., 2005; Bull and Lauring, 2014; Gandon et al., 2016). A classical explanation for pathogen virulence involves trade-offs with other pathogen life-history traits. If certain components of pathogen fitness, such as a high transmission rate or a long duration of transmission, necessarily require that the pathogen incidentally harm its host then virulence is expected to evolve (Frank, 1996). A now classical way to develop specific predictions from this hypothesis is to adopt the Adaptive Dynamics formalism (Geritz et al., 1998; Metz et al., 1992; Dieckmann et al., 2005; Frank, 1996). This approach relies on the assumption that the mutation rate is small so that the epidemiological dynamics occur on a faster timescale than the evolutionary dynamics (Anderson et al., 1992; Frank, 1996; Alizon et al., 2009; Cressler et al., 2016). Under simple epidemiological assumptions (no co-infections with different genotypes) the evolutionarily stable level of virulence maximizes the basic reproduction ratio *R*_0_ of the pathogen (but see *e.g.*, Nowak and May (1994); van Baalen and Sabelis (1995) for more complex epidemiological scenarios).

Adaptive Dynamics models allow one to predict long-term evolution but they tell us little about what we should expect to observe if epidemiological and evolutionary processes occur on a similar timescale. Novel theoretical approaches have therefore been developed to address this issue (Lenski and May, 1994; Frank, 1996; Day and Proulx, 2004; Day and Gandon, 2006; Bull and Ebert, 2008). These studies have revealed that, in addition to tradeoffs, the nature of the epidemiological dynamics (e.g., epidemic spread versus endemic disease) can also dictate the type of pathogen that will evolve. For example, pathogens with relatively high virulence can be selected for during epidemic disease spread whereas pathogens with a lower virulence may outcompete such high virulence strains in endemic diseases (Lenski and May, 1994; Frank, 1996; Day and Proulx, 2004; Berngruber et al., 2013).

The above-mentioned theory allows one to determine the level of virulence expected to evolve under a broad range of epidemiological scenarios but it still suffers from the fundamental shortcoming of being a deterministic theory. Pathogen population size, however, can be very small (e.g. at the onset of an epidemic or after a vaccination campaign) and demographic stochasticity is likely to affect both the epidemiological and evolutionary dynamics of the disease. If all that such stochasticity did was to introduce random noise then the predictions of deterministic theory would likely suffice. However, several recent studies have demonstrated that this is not the case. For example, Kogan et al. (2014) and Humplik et al. (2014) each used different theoretical approaches to demonstrate that finite population size tends to select for lower virulence and transmission. Keeling (Likewise, Read and 2007) analyzed the effect of finite population size in a complex epidemiological model with unstable epidemiological dynamics and showed that finite population size could induce an evolutionary instability that may either lead to selection for very high or very low transmission.

Taken together, the existing literature presents a complex picture of the factors that drive virulence evolution and it remains unclear how all of these factors are related to one another and how they might interact. In this paper we develop a very general theory of pathogen evolution that can be used to examine virulence evolution when all of the above-mentioned factors are at play. First, we use an individual based description of the epidemiological process to derive a stochastic description of the evolutionary epidemiology dynamics of the pathogen. This theoretical framework is used to pinpoint the effect of finite population size on the interplay between epidemiology and evolution. Second, we analyze this model under the realistic assumption that the rate of mutation is small so that pathogen evolution can be approximated by a sequence of mutation fixations. We derive the probability of fixation of a mutant pathogen under both weak and strong selection regimes, and for different epidemiological scenarios. Third, we use this theoretical framework to derive the stationary distribution of pathogen virulence resulting from the balance between mutation, selection and genetic drift. This yields new predictions regarding the effect of the shape of pathogen fitness landscape, the size of the population and sources of host heterogeneities on long-term evolution of the pathogen. As the question of virulence evolution can be viewed as a specific example of the more general notion of life history evolution (Stearns, 1992; Roff, 2002) our results should be directly applicable to other life history traits and other organisms as well.

## 2 Model

We use a classical SIR epidemiological model where hosts can either be susceptible, infected or recovered. The number of each of these types of hosts is denoted by *N_S_, N_I_*, and *N_R_* respectively. Because we are interested in the effect of demographic stochasticity the model is derived from a microscopic description of all the events that may occur in a finite host population of total size *N_T_ = N_S_ + N_I_ + N_R_* (the derivation of the model is detailed in Supplementary Information). It will be useful to explicitly specify the size of the habitat in which the population lives (e.g., the area of the habitat) and so we denote this by the parameter *n*.

We use λ to denote the rate at which new susceptible hosts enter the population per *unit area* and therefore the total rate is given by *λn*. We focus on the case of frequency-dependent transmission; i.e., new infections occur at rate 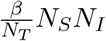 where *β* is a constant quantifying the combined effects of contact rate among individuals and the probability of pathogen transmission given an appropriate contact occurs. Note, however, that other forms of transmission (*e.g.*, density dependent transmission,(McCallum et al., 2001)) yield qualitatively similar results (Parsons, 2012). For simplicity we also assume that already infected hosts cannot be reinfected by another pathogen strain (i.e., no co-infections). All hosts are assumed to suffer a constant per capita death rate of *δ* and infected hosts die at per capita rate *α* and they recover at per capita rate *γ*. Finally, to study pathogen evolution we need to introduce genetic variation in the parasite population. Therefore we consider *d* pathogen strains which differ in transmission rate *β_i_* and virulence *α_i_*, with *i* ∈ {1,…, *d*}. Likewise we use the subscripted variable *N_I_i__* to denote the number of hosts infected with strain *i*.

The dynamical system resulting from the above assumptions is a continuous-time Markov process tracking the number of individuals of each type of host. To progress in the analysis we use a diffusion approximation and work with host densities defined as *S* = *N_S_*/*n*, *I_i_* = *N_I_i__*/*n* and *N* = *N_T_/n* and we define the total density of infected hosts as 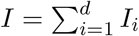. When n is sufficiently large these variables can be approximated using a continuous state space and so this model can be described by a system of stochastic differential equations (see Supplementary Information, §3).

### 2.1 Deterministic evolution

In the limit where the habitat size (and thus the host population size) gets large, demographic stochasticity becomes unimportant and the epidemiological dynamics are given by the following system of ordinary differential equations:

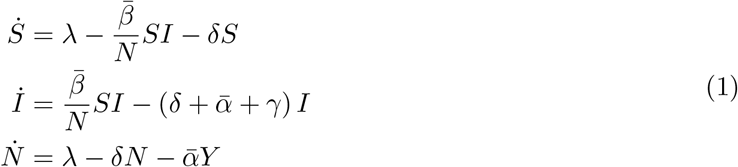

The bars above *α, β* and *γ* refer to the mean of the transmission and the virulence distributions of the pathogen population (i.e. 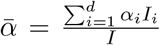). In the absence of the pathogen the density of hosts equilibrates at 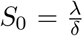. A monomorphic pathogen population (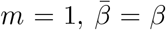 and 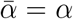) is able to invade this equilibrium if its basic reproduction ratio is 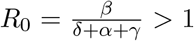. If this condition is fulfilled the system reaches an endemic equilibrium where 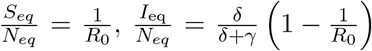 and 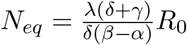.

When several strains are present in the population the evolutionary dynamics of the pathogen can be tracked with (Day and Gandon, 2006, 2007):

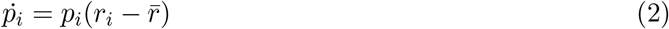

where 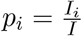 is the frequency of pathogen *i*. The quantity 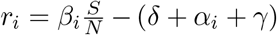 is the instantaneous per capita growth rate of strain *i* and 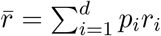 is the average per capita growth rate of the pathogen population. When *m* = 2 only two strains are competing (a wild-type, strain 1, and a mutant, strain 2) the change in frequency *p*_2_ of the mutant strain is given by:

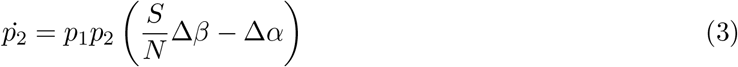

where Δ*β* = *β*_2_ – *β*_1_ and Δ*α* = *α*_2_ – *α*_1_ are the effects of the mutation on transmission and virulence, respectively.

The above formalization can be used to understand the evolution of pathogen life-history under different scenarios. First, under the classical Adaptive Dynamics assumption that mutation rate is very small one may use a separation of time scales where the epidemiological dynamics reach an endemic equilibrium (set by the resident pathogen, strain 1) before the introduction of a new variant (strain 2) by mutation. In this case evolution favours the strain with the highest basic reproduction ratio: 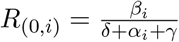. In other words, evolution favours strains with higher transmission rates and lower virulence. According to the tradeoff hypothesis, however, transmission and virulence cannot evolve independently. For example, the within-host growth rate of pathogens is likely to affect both traits and result in a functional trade-off between transmission and virulence (Anderson et al., 1992; Frank, 1996; Alizon et al., 2009; Cressler et al., 2016). Under this assumption equation (3) can be used to predict the evolutionary stable virulence strategy (Figure 1). The above model can also be used to predict virulence evolution when the evolutionary and epidemiological dynamics occur on a similar time scale (Day and Gandon, 2006, 2007; Gandon and Day, 2007). For instance, these models can be used to understand virulence evolution during an epidemic (Lenski and May, 1994; Frank, 1996; Day and Proulx, 2004; Berngruber et al., 2013). In this case, a pathogen strain *i* with a lower *R*_0_ may outcompete other strains if its instantaneous growth rate, *r_i_*, is higher.

**Figure 1:**
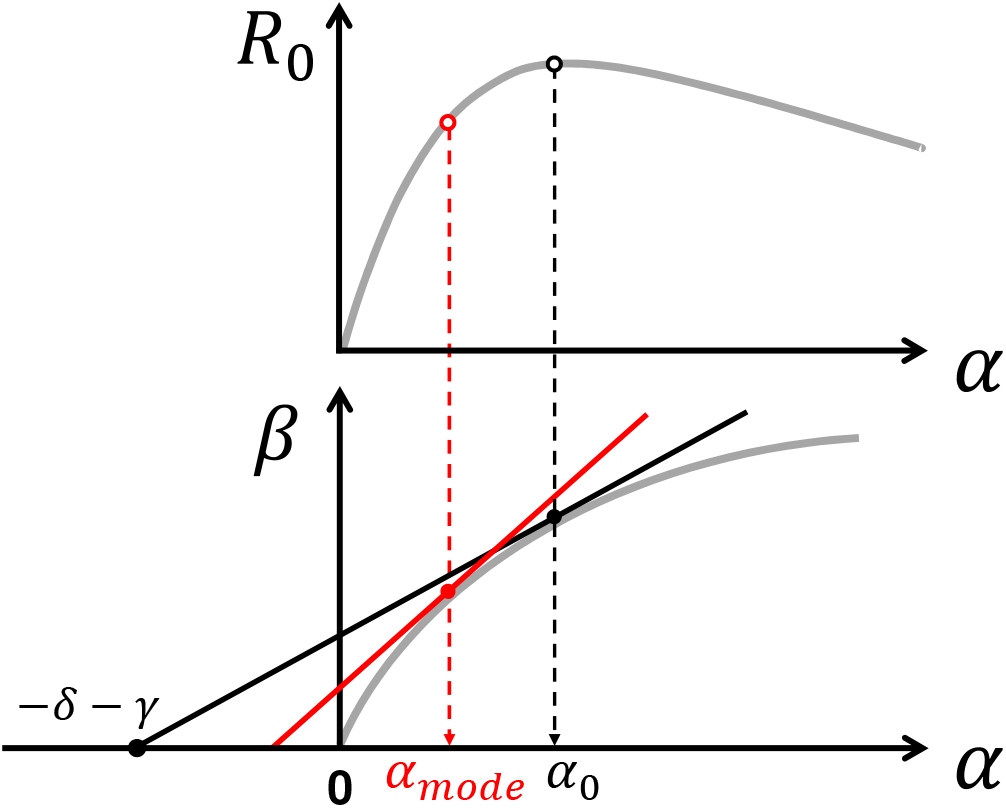
Schematic representation of the effect of finite population size on the evolution of pathogen virulence. The grey line in the top figure represents the effect of pathogen virulence, *α*, on *R*_0_ (for an asymmetric fitness function). The grey line in the bottom figure represents the effect of pathogen virulence, *α*, on pathogen transmission, *β*. In the deterministic version of our model the marginal value theorem can be used to find the evolutionary stable (ES) pathogen virulence, *α*_0_ (dashed black arrow). In this model ES virulence maximizes *R*_0_ in the absence of demographic stochasticity. Finite population size modifies selection and favours pathogen strategies with lower virulence (see equation (10)). The mode of the stationary distribution of pathogen virulence is indicated by a dashed red arrow, *α*_mode_ (see equation (13)). This geometrical construction indicates that finite population size is expected to favour *slower* strains even if they have a lower *R*_0_.

### 2.2 Stochastic evolution

Finite population size introduces demographic stochasticity and the epidemiological dynamics can be described by the following system of (Itô) stochastic differential equations:

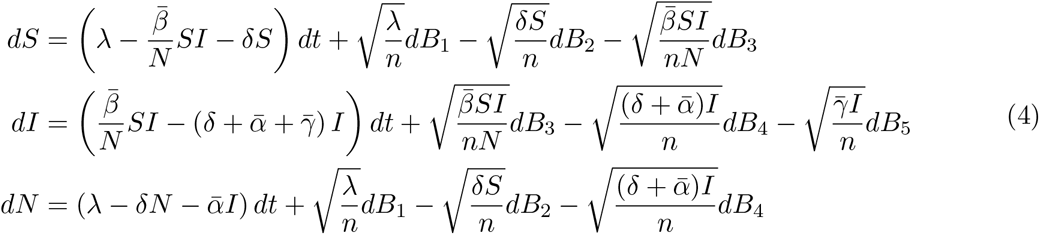

where *B*_1_,…, *B*_5_ are independent Brownian motions. As expected, when *n* → ∞ this set of stochastic differential equations reduces to the deterministic equations in (1).

In finite populations the pathogen, and indeed the host population itself, are destined to extinction with probability 1. The time it takes for this to occur, however, depends critically on the parameter values. For example, in a monomorphic pathogen population (i.e., *m* = 1), if *R*_0_ is larger than one the size of the pathogen population reaches a quasi-stationary distribution which is approximately normal. The mean of this distribution is of order *n* and its standard deviation is of order 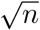 (Nåsell, 2001, 2007). The extinction time from the quasi-stationary distribution increases exponentially with *n* (Barbour, 1976; Nåsell, 2001, 2007) and so, in the remainder of the paper we will assume that *n* is large enough so that we can focus on the dynamics conditional on non-extinction.

As in the deterministic case, one can study evolutionary dynamics by focusing on the change in strain frequencies. We obtain a stochastic differential equation analogous to (2) (see Supplementary Information, §4):

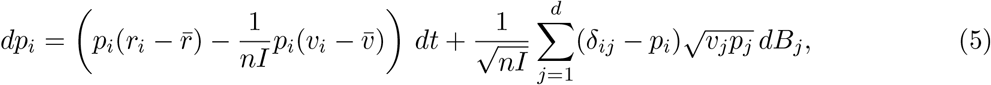

where 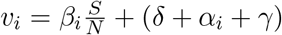 is the variance in the growth rate of strain *i* (while *r_i_* is the mean) and 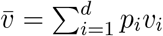 is the average variance in growth rate of the pathogen population. The first term in equation (5) is analogous to (2). The second term shows that finite population size (*i.e*., when pathogen population size, as measured by the total density of infected hosts, nI is not too large) can affect the direction of evolution. In contrast with the deterministic model, the evolutionary dynamics are not driven exclusively by the expected growth rate *r_i_* but also by a minimization of the variance. This effect is akin to bet-hedging theory stating that a mutant strategy with lower variance in reproduction may outcompete a resident strategy with a higher average instantaneous growth rate (Gillespie, 1974; Frank and Slatkin, 1990). To better understand this effect it is particularly insightful to examine the case *m* = 2 when only two strains are competing and the change in frequency *p*_2_ of the mutant strain is given by:

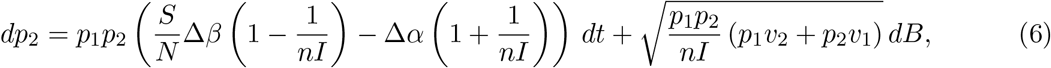

The first term (the drift term) in equation (6) is similar to (3) except for the 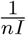 terms. Those terms are due to the fact that a transmission (or a death) event of the mutant is associated with a change in the number of mutants as well as an increase (decrease) of the total pathogen population size by one individual. This concomitant variation of pathogen population size affects the effective change of the mutant frequency (relative to the change expected under the deterministic model where population size are assumed to be infinite). This effect decreases the benefit associated with higher transmission and increases the cost of virulence. In the long-term this effect (the drift term in (5)) is thus expected to select for lower virulence. But this long term evolutionary outcome cannot be described by an evolutionary stable state because demographic stochasticity is also expected to generate noise (the diffusion term in (5)). Indeed, this stochasticity (*i.e.*, genetic drift) may lead to the invasion and fixation of strains with lower per capita growth rates. In the following we fully characterize this complex evolutionary outcome with the stationary distribution of pathogen virulence under different epidemiological scenarios.

## 3 Results

The above theoretical framework embodied by the stochastic differential equations (4) and (5) subsume the deterministic model and can be used to study the interplay of all the relevant factors affecting virulence evolution. In the following we will assume that pathogen mutation is rare so that evolution can be described, as in classical Adaptive Dynamics, as a chain of fixation of new pathogen mutations. In contrast with Adaptive Dynamics, however, demographic stochasticity may allow deleterious mutations to go to fixation. The analysis of the effect of finite population size requires specific ways to quantify the stochastic fate of a genotype (Proulx and Day, 2002). To determine the fate of a new mutation we need to compute the probability of fixation of a mutant pathogen in a resident population. In the absence of selection the fixation probability of a mutant allele depends only on the demography of the population. When the size of the population is fixed and equal to *N* the fixation probability of a neutral allele is 1/*N*. When the fixation probability of a mutant is higher than neutral it indicates that the mutant is selectively favoured. This is particularly useful in many complex situations where the interplay between selection and genetic drift are difficult to disentangle like time varying demography (Otto and Whitlock, 1997; Lambert, 2006) or spatial structure (Rousset, 2004). In our model, the difficulty arises from (i) the stochastic demography of the pathogen population and (ii) the fact that pathogen life-history traits feed-back on the epidemiological dynamics and thus on the intensity of genetic drift.

### 3.1 Stationary distribution of pathogen virulence at equilibrium

Here we assume, as in the Adaptive Dynamics framework, that the pathogen mutation rate μ is so low that the mutant pathogen (strain 2) arises when the resident population (strain 1) has reached a stationary equilibrium *nI_eq_* (*i.e*., close to the endemic equilibrium derived in the deterministic model). The *R*_0_ of the two strains may be written in the following way: *R*_0,2_ = *R*_0,1_ (1 + *s*) where s measures the magnitude of selection.

When selection is strong (*i.e., s* ≫ *n*) the probability of fixation of the mutant when *N*_*I*_2__(0) mutants are introduced into a resident population at equilibrium is (see Supplementary Information, §5.2):

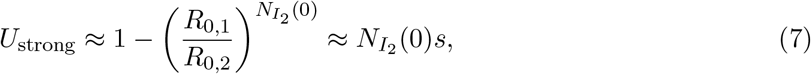

which may be obtained by approximating the invading strain by a branching process (see Supplementary Information, §7.2 for a rigorous justification). When the mutant and the resident have similar values of *R*_0_ (*i.e., s* is of order 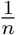) selection is weak, and the derivation of the probability of fixation is a much more difficult problem. The classical population genetics approach under the assumption that population size is fixed (or is characterized by a deterministic trajectory independent of mutant frequency) is to use the diffusion equation of mutant frequency to derive the probability of fixation (Otto and Whitlock, 1997; Lambert, 2006). But in our model, equation (3) is not autonomous and is coupled with the epidemiological dynamics. To derive the probability of fixation we use a separation of time scale argument to reduce the dimension of the system (see (Parsons and Rogers, 2017) for a discussion of the approach). Indeed, if selection is weak the deterministic component of the model sends the system rapidly to the endemic equilibrium. At this point, it is possible to approximate the change in frequency of the mutant by tracking the dynamics of the projection of the mutant frequency on this manifold (see Supplementary Information, §5.3). This one dimensional system can then be used to derive the probability of fixation under weak selection. A first order approximation in *s* and *σ* is:

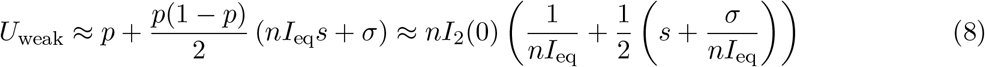

where *p* = *I*_2_(0)/*I_eq_* and 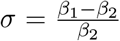. The first term in (8) is the probability of fixation of a single neutral mutation introduced in a pathogen population at the endemic equilibrium *nI_eq_*. The second term takes into account the effect due to selection. First, selection may be driven by differences in *R*_0_. Second, even if strains have identical *R*_0_ (*i.e., s* = 0) selection may be driven by *σ* which measures the difference in transmission rate. Note, however, that the effect of s rapidly overwhelms the effect of *σ* as pathogen population size *nI_eq_* becomes large (unless s is of order 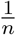). The probability of fixation given in (8) confirms that evolution tends to push towards higher basic reproductive ratio but when the population size is small other forces may affect the evolutionary outcome. In particular, when *nI_eq_* is small, strains with lower *R*_0_ can reach fixation. Figure 2 shows the result of stochastic simulations that confirm the approximations (7) and (8) under different epidemiological scenarios.

**Figure 2:**
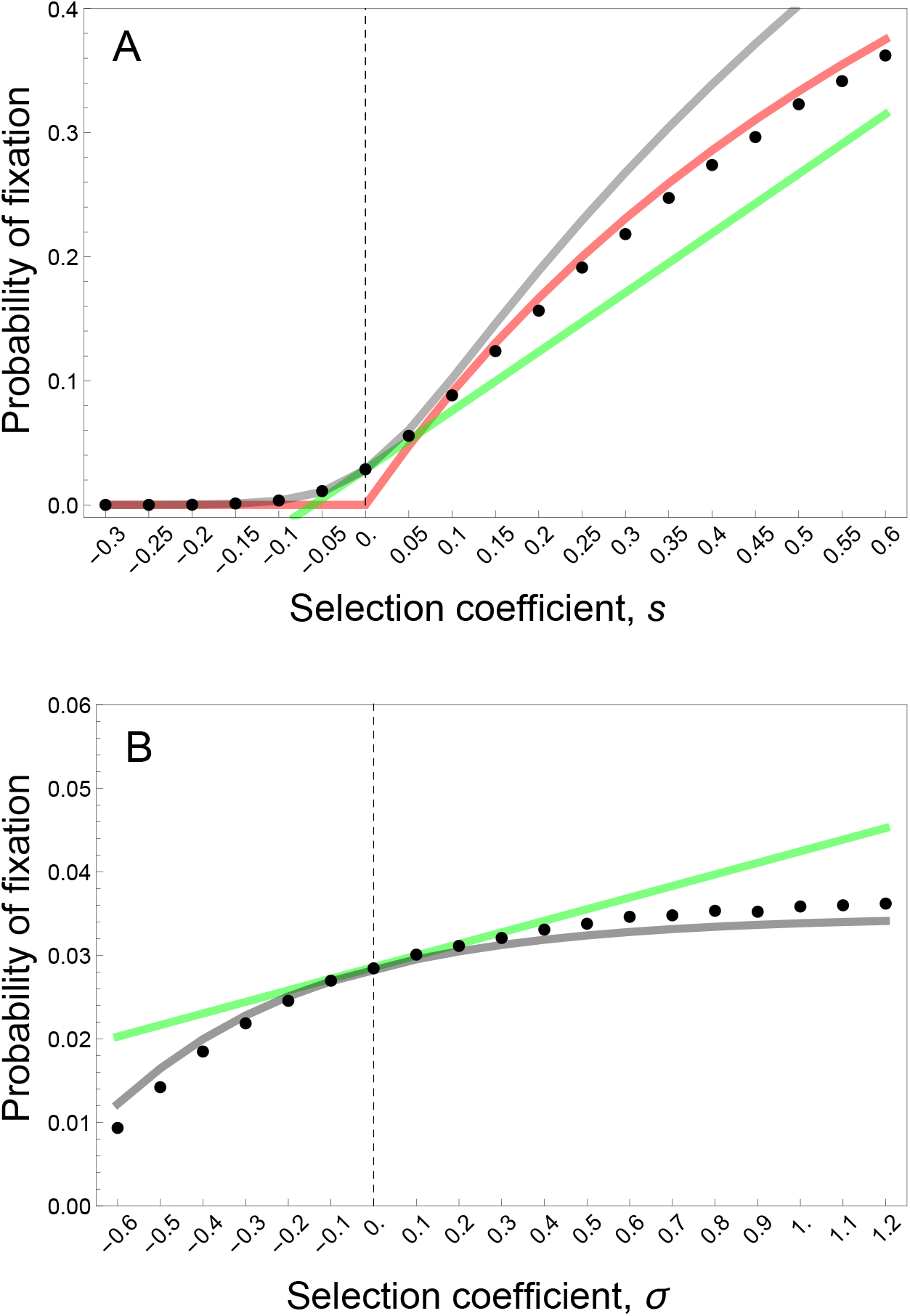
Probability of fixation for (A) different values of *s* (strong selection effect) and (B) different values of *σ* (weak selection effect). Simulation results are indicated with a dot, weak selection approximation is indicated with a gray line and its linear approximation (equation (8)) is indicated with a green line, the strong selection approximation is indicated with a red line (equation (7)). Parameter values of the resident population: *n* = 100, *R*_0_ = 4, *δ* = 1, *α* = 3, *γ* = 1, *λ* = 2, *β*_1_ = 20. For the simulation a single mutant (an individual host infected with a mutant pathogen) is introduced at the endemic equilibrium set by the resident pathogen: *S*_eq_ = 24 and *I*_eq_ = 35. 10^6^ simulations are realized for each parameter values and we plot the proportion of the simulations where the mutant goes to fixation.

Even though the probability of fixation helps understand the interplay between selection and genetic drift it does not account for any differences in the time to fixation and it is often difficult to measure this probability experimentally as well (but see Gifford et al. (2012)). What may be more accessible is a characterization of the phenotypic state of the population across different points in time (or in space among replicate populations) - that is, the stationary distribution of the virulence phenotype of the pathogen under the action of mutation, selection and genetic drift (Champagnat and Lambert, 2007; Lehmann, 2012; Debarre and Otto, 2016) (Figure 3).

**Figure 3:**
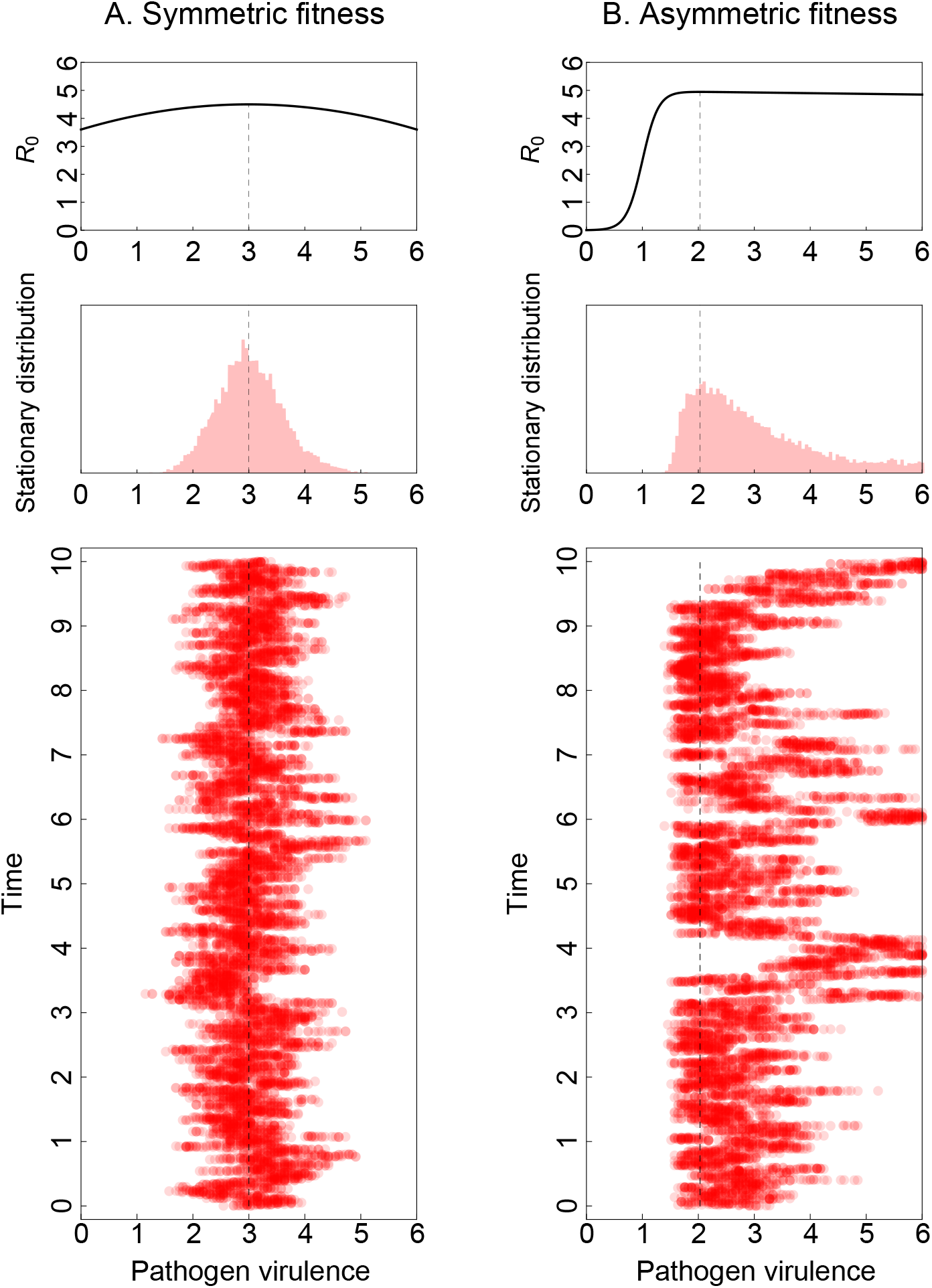
Dynamics of pathogen virulence across time (one time unit on the graph is 10^7^ time steps in the simulation, and a) and stationary distribution of pathogen virulence for two different fitness landscapes: (A) Symmetric fitness landscape with *β*(*α*) = (*δ + γ + α*)*R*_0,max_ (1 – *w*(*α*_0_ – *α*)^2^), *R*_0,max_ = 4.5 and *α*_0_ = 3, (B) Asymmetric fitness landscape with *β*(*α*) = 5(1 – 0.005α)/(1 + exp(7(1 – *α*)). The dashed vertical line indicates the position of *α*_0_. Other parameter values: *n* = 200, *δ* = 1, *α* = 3, 7 = 1, *λ* = 2, *μ* = 0.001.

To derive the stationary distribution of pathogen virulence we first need to impose a trade-off between virulence and contact rate, setting *β* = *β*(*α*), and introduce the mutation kernel *K*(*α_m_, α*), the probability distribution of mutants with strategy *α_m_* from a monomorphic population with strategy *α*. Here we assume that this distribution is Gaussian with a mean equal to the current resident trait value and variance *v*. Under the assumption that the mutation rate *μ* remains small, pathogen polymorphism is limited to the transient period between the introduction of a mutant and a fixation. The probability of fixation (8) accurately describes the direction of evolution and the evolution of pathogen virulence can then be described by the following Fokker-Planck diffusion equation (see Supplementary Information, §6):

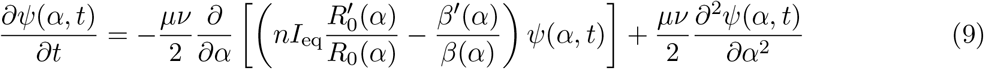

where *ψ*(*α, t*) is the distribution of pathogen virulence and’ indicates the derivative with respect to α. The drift term of the above equation indicates that deterministic evolution tends to maximize the basic reproduction ratio while finite population size tends to select for lower transmission. Under the classical assumption that pathogen transmission and pathogen virulence are linked by a genetic trade-off one can ask what the level of pathogen virulence is where the drift coefficient is zero. This trait value corresponds to the mode of the stationary distribution of pathogen virulence and is given by the following condition (see Supplementary Information Equation S.45):

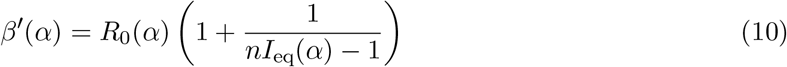

When the pathogen population is very large (i.e.,*n* → ∞) we recover the marginal value theorem while finite population size increases the slope *β*′(*α*) and reduces the mode of the stationary distribution (see Figure 1). Thus, for a broad range of transmission-virulence trade-off functions, finite population size is expected to decrease virulence and transmission rates. In other words, pathogen avirulence may be viewed as a bet-hedging strategy because even if it reduces the instantaneous growth rate *r_i_*, the reduced variance in growth rate *υ_i_* is adaptive in finite population size.

Let us now consider the limiting case when all the pathogen strains have the same R_0_. This corresponds to a very special case where the fitness landscape is flat. The deterministic model predicts that pathogen life-history variation is neutral near the endemic equilibrium (see (2)). The probability of fixation (8) shows, however, that selection is *quasi-neutral* and favours pathogens with lower transmission and virulence rates (Parsons and Quince, 2007; Parsons, 2012; Kogan et al., 2014; Humplik et al., 2014). The stationary distribution results from the balance between selection (pushing towards lower values of pathogen traits) and mutation (reintroducing variation). If we focus on virulence and allow variation between a minimal value *α*_min_ and a maximal value *α*_max_ the stationary distribution is (see Supplementary Information Equation S.39):

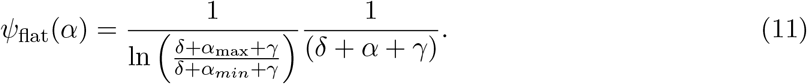

It is worth noting that this distribution is independent of the pathogen population size. Indeed, near the endemic equilibrium and when pathogens have the same *R*_0_ the probability of fixation (8) is independent of pathogen population size. So this prediction holds even in very large pathogen populations. The time to fixation may, however, be considerably longer in large populations and the assumption that polymorphism is always reduced to the resident and a single mutant may not always hold as pathogen population increases. Yet, stochastic simulations confirm that (11) correctly predicts the stationary distribution, which is relatively insensitive to pathogen population size but varies with *δ* + *γ* (Figure 4A).

**Figure 4:**
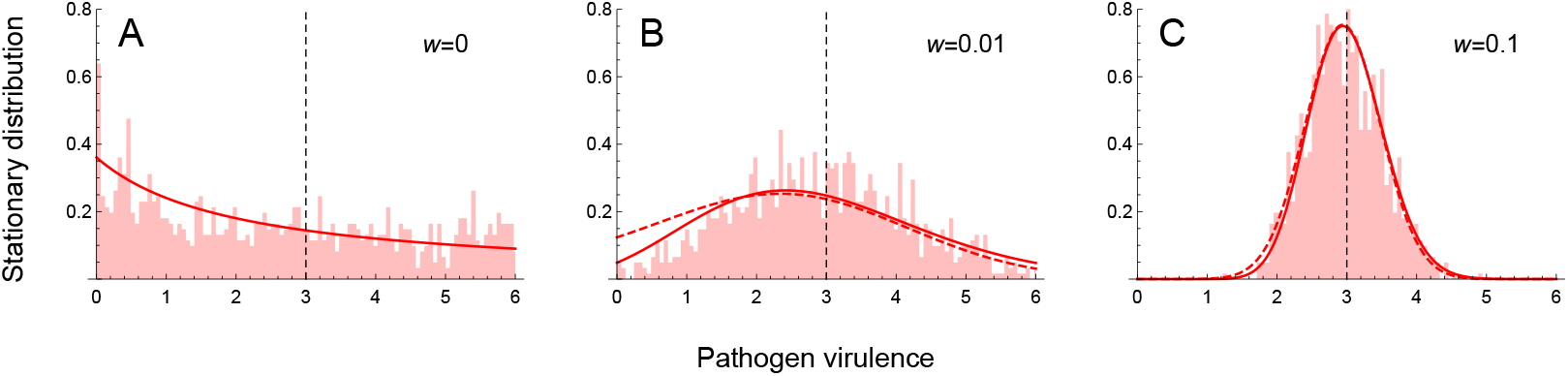
Stationary distribution for symmetric fitness landscapes with increasing strength of selection around the optimum with *β*(*α*) = (*δ* + *γ* + *α*)*R*_0,max_ (1 – *w*(*α*_0_ – *α*)^2^) and *α*_0_ = 3 for three different values of *w*: (A) *w* = 0, (B) 0.01 and (C) 0.1. Note that when *w* = 0 the fitness landscape is flat. The light red histogram indicates results of a stochastic simulation. The red line indicates the stationary distribution of the diffusion approximation (the dashed line indicates the approximation of this distribution, see (12)). The dashed vertical line indicates the position of *α*_0_. Parameter values: *n* = 200, *R*_0,max_ = 4, *d* = 1, *α* = 3, *γ* = 1, *λ* = 2, *μ* = 0.001.

Second, we consider a general fitness landscape with a single maximum. It is possible to derive a good approximation for the stationary distribution (see S.44 in the Supplementary Information):

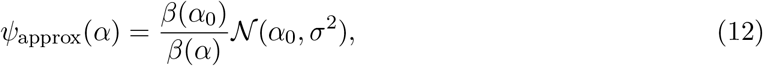

where *α*_1_ is the virulence that maximizes *R*_0_, *I_eq_*(*α*_0_) is the expected number of infected individuals at the endemic equilibrium when the virulence is *α*_0_ and 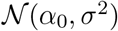 is the Gaussian distribution with mean *α*_0_ and variance 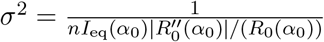. We thus see the effect of the demography is to bias the Gaussian, putting more weight on values of the virulence below *α*_0_; this becomes more clear when we consider the mode and mean of the (true) stationary distribution (see §6.3 in the Supplementary Information):

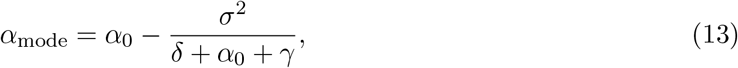

and

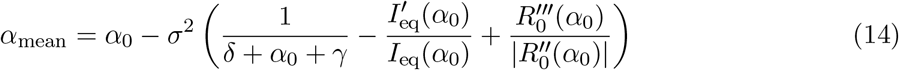

respectively. These results indicate that, as expected from the simple optimization approach used above in (10) and illustrated in Figures 3 and 4, lower pathogen population size tends to decrease pathogen virulence. Yet, the above derivation of the stationary distribution goes far beyond this optimization criterion. First, it predicts accurately the mode of the stationary distribution. In particular it shows that the shape of the fitness landscape may affect the mode of the stationary distribution. The skew of the fitness landscape can have huge effects on the stationary distribution (Figure 3). A positive skew leads to a higher mean virulence and may thus counteract the effect of small pathogen population. In other words, whether demographic stochasticity favours lower of higher virulence depends also on the shape of the fitness landscape. Second, our analysis predicts the amount of variation one may expect to see around this mode. Unlike the criteria used to derive a single optimal strategy our approach predicts accurately the expected variation around this mode (Figures 3 and 4). Note that the population remains monomorphic most of the time (because mutation is assumed to be small) but the variance of the stationary distribution refers to the distribution of phenotypes explored through time (or through space if stochastic evolution is taking place in multiple isolated populations).

### 3.2 Pathogen evolution after vaccination

The above analysis relies on the assumption that the epidemiological dynamics are much faster than evolutionary dynamics so that the pathogen population is always at its endemic equilibrium. The present framework can also be used to explore the fate of a mutant pathogen away from this endemic equilibrium. For instance, right after the start of a vaccination campaign the availability of susceptible hosts is going to drop rapidly if the vaccination coverage *f* is high and if the efficacy of the vaccine is large. This epidemiological perturbation has major consequences on the probability of fixation of a pathogen mutant. Looking at invasion out of endemic equilibrium, we show that a larger vaccination coverage *f* decreases the probability of fixation of all pathogen mutants (see Supplementary Information §5.2.3) but the probability of fixation of strains with low virulence and low transmission rates are less affected than more virulent and transmissible strains. In other words, strains with low turn over rates (*slower* strains) are less likely to be driven to extinction during the drop of the pathogen population size. This extends classical results of populations genetics (Otto and Whitlock, 1997; Lambert, 2006) to situations where the mutations are acting on life history traits and feed back on population dynamics. *Faster* strains are selected for during epidemics while *slower* strains are favoured when the pathogen population is reduced (e.g. after a public health intervention). This is also consistent with the analysis of pathogen evolution based on deterministic models which showed that the direction of selection depends on the epidemiological state of the population (Lenski and May, 1994; Frank, 1996; Day and Proulx, 2004; Lambert, 2006; Day and Gandon, 2006, 2007; Berngruber et al., 2013).

Vaccination is also meant to induce long-term modifications of the host population. In particular, artificial immunization introduces a heterogeneity between vaccinated and unvaccinated hosts. If the vaccine is perfect, vaccination will act on the epidemiology and will reduce the endemic equilibrium. In a deterministic version of this model, such a perfect vaccine is expected to have no consequences on long-term pathogen evolution (Gandon et al., 2001; Gandon and Day, 2007). In contrast, when host population size is finite, vaccination is expected to magnify the influence of demographic stochasticity and, as discussed above, to select for lower pathogen virulence and to increase the variance of the stationary distribution (Figure 5). It is also interesting to consider an alternative scenario where vaccinated hosts can be infected but cannot transmit the pathogen. In this case, two types of infected hosts are coexisting: good-quality (naïve) hosts and bad-quality (vaccinated) hosts. In this situation, the amount of demographic stochasticity is not governed by the whole pathogen population size but by the size of the population of infected hosts that actually contribute to transmission. In fact all the results derived above apply in this scenario provided that the equilibrium pathogen population size is replaced by the *effective* pathogen size: *nI_e_* = (1 – *f*)*nI*_eq_. This example illustrates that even for large pathogen population sizes the effect of demographic stochasticity can be important if the *effective* pathogen size is small. When there is variation in reproductive value among individuals the *effective* population size that governs the amount of genetic drift may be substantially lower than the *actual* size of the population (Crow and Kimura, 1970). In the context of pathogen evolution this heterogeneity in reproductive value may be driven by variations in infectiousness among hosts. This variation can be induced by public-health interventions (e.g. transmission-blocking vaccines) but it emerges naturally from complex behavioural and/or physiological differences among hosts. For instance, evidence for *superspreading* events where certain individuals can infect unusually large numbers of secondary cases have been found in many human pathogens (Woolhouse et al., 1997; Lloyd-Smith et al., 2005). This heterogeneity is expected to reduce the effective population size and to magnify the effects of demographic stochasticity discussed above. But other factors like temporal fluctuations in population size are also known to modulate the intensity of genetic drift and may affect effective population size (Crow and Kimura, 1970).

**Figure 5:**
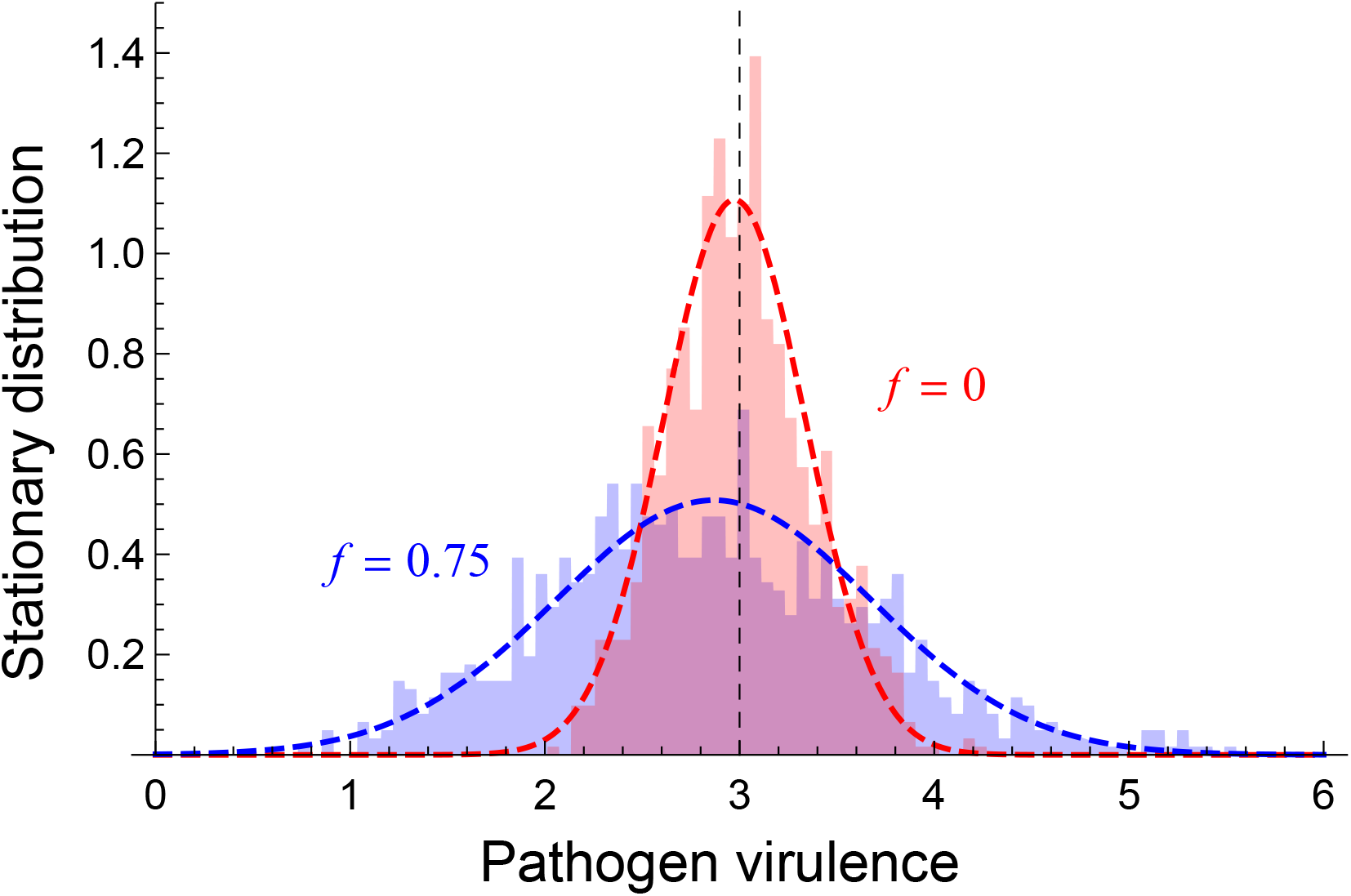
Stationary distribution for symmetric fitness landscapes with increasing vaccination coverage (*f* = 0 in red and *f* = 0.75 in blue) with *β*(*α*) = (*δ* + *γ + α*)*R*_0,max_ (1 – *w*(*α*_0_ – *α*)^2^) and *α*_0_ = 3. The histogram indicates results of a stochastic simulation. The blue and red dashed lines indicate the approximation of the stationary distribution given in (12) with or without vaccination, respectively. The dashed vertical line indicates the position of *α*_0_. Parameter values: *n* = 1000, *R*_0,max_ = 10, *δ* = 1, *α* = 3, *γ* = 1, *λ* = 2, *μ* = 0.01, *w* = 0.1

## 4 Discussion

Evolutionary theory has led to the development of different mathematical tools for studying phenotypic evolution in a broad diversity of ecological scenarios (Parker and Smith, 1990; Roff, 1993; Otto and Day, 2007). For instance, Adaptive Dynamics is a powerful theoretical framework to study life-history evolution when mutation is assumed to be rare so that demographic and evolutionary processes can be decoupled (Geritz et al., 1998; Metz et al., 1992). This analysis yields evolutionarily stable life-history strategies and captures the ultimate outcome of evolution. But this approach relies on the assumption that population size is infinite and that evolution is deterministic. Finite population size, however, can also affect evolutionary trajectories. In particular, even the fittest genotype can be invaded by a deleterious mutant when population size is reduced. This leads to the collapse of the concept of evolutionarily stable strategy. Here we develop and apply Stochastic Adaptive Dynamics (SAD) (Champagnat and Lambert, 2007; Otto and Day, 2007; Debarre and Otto, 2016), a new theoretical framework where the evolutionary outcome of life history evolution is studied through the derivation of the stationary distribution of the phenotype under mutation-selection-drift equilibrium. Under the assumption that mutation rate is small, the equilibrium distribution can be derived from a diffusion approximation. In contrast with previous population genetics models, the present framework also allows life-history evolution to affect population size and, consequently, the amount of demographic stochasticity. In other words, this framework retains key features of Adaptive Dynamics but relaxes a major assumption by allowing genetic drift to affect the evolutionary outcome (see also Waxman and Gavrilets, 2005, p.1149). As such, our SAD framework is an important step towards a better integration between Adaptive Dynamics and classical population genetics.

We show that finite population size induces a selective pressure towards strains with lower variance in growth rate (but see also Gillespie (1974); Lambert (2006)). A simple way to understand this effect is to compare the fate of two strains with the same R0 but with different life-history strategies. The *fast* strain is very transmissible but has a short duration of infection (e.g., because of high virulence or high clearance rate). The *slow* strain has a long duration of infection but has a small transmission rate. Since the two strains have the same R0 Adaptive Dynamics predicts that these two strains should coexist. With finite population size, however, the fast strain has a higher probability to go extinct simply because more events happen per unit of time. As in Aesop’s Fable “Slow and steady wins the race” because the fast strain will reach extinction sooner than the slow strain. Previous studies (Parsons, 2012; Kogan et al., 2014; Humplik et al., 2014) pointed out the influence of finite population size on the direction of virulence evolution but they focused mainly on the quasi-neutral case where all the strains have the same *R*_0_. Humplik et al. (2014) did look at scenarios where strains have different R0 but without a derivation of the stationary distribution at mutation-selection-drift equilibrium. We believe that this stationary distribution is key to explore the interaction between finite population size and phenotypic evolution. This distribution yields testable predictions on the mean as well as other moments of the phenotypic distribution.

The approximation (12) shows that this distribution is moulded by two main parameters: (i) the pathogen fitness landscape, and (ii) the effective size of the pathogen population. First, the fitness landscape at the endemic equilibrium can be derived from (5) and depends mainly on the way *R*_0_ varies with pathogen life history traits. Under the classical transmission-virulence assumption R0 is maximized for some intermediate virulence. But the shape of the trade-off also affects the shape of the fitness landscape and in particular its symmetry. Second, the effective size *nl_e_* of the pathogen population size depends mainly on the pathogen population size *nl_eq_* at the endemic equilibrium but other factors may reduce the effective pathogen population size as well. For instance, variance in transmission among infected hosts is likely to reduce *nl_e_* below *n*/_eq_. One source of heterogeneity in transmissibility may be induced by public-health interventions (e.g., vaccination, drug treatments), but intrinsic behavioural or immunological heterogeneities among hosts may induce superspreading transmission routes as well (Woolhouse et al., 1997; Lloyd-Smith et al., 2005).

When the fitness landscape of the pathogen is symmetric, reducing the effective population size increases the variance of the stationary distribution but decreases also the mean (and the mode) of this distribution. This effect results from the selection for a reduction of the variance identified in (5). This is the effect that emerges in the quasi-neutral case. When the fitness landscape is flat this may lead to an important bias towards lower virulence (Figure 4). When the fitness landscape of the pathogen is asymmetric the skewness of the fitness landscape can affect the mean of the stationary distribution when the effective population size *nI_e_* of the pathogen is reduced. More specifically negative (positive) skewness reduces (increases) the mean of the stationary distribution. It is interesting to note that classical functions used to model the trade-off between virulence and transmission tend to generate positive skewness in the fitness landscape (van Baalen and Sabelis, 1995; Frank, 1996; Alizon et al., 2009). The asymmetry of these fitness functions may thus counteract the effects of stochasticity per se identified in symmetric fitness landscapes. In other words, predictions on the stochastic evolutionary outcome are sensitive to the shape on genetic constraints acting on different pathogen life-history traits. This result is very similar to the deterministic effects discussed in (Urban et al., 2013) on the influence of asymmetric fitness landscapes on phenotypic evolution. Note, however, that the effect analyzed by Urban et al (2013) is driven by environmental effects on phenotypes. In our model, we did not assume any environmental effects and a given genotype is assumed to produce a single phenotype.

We focused our analysis on the stationary distribution at the endemic equilibrium of this classical SIR model. But we also explore the effect of demographic stochasticity on the transient evolutionary dynamics away from the endemic equilibrium. For instance we recover a classical population genetics result (Otto and Whitlock, 1997; Lambert, 2006) that the probability of fixation of adaptive mutations (*i.e.*, with 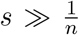) is increased during epidemics. Beyond the effect of *s* (*i.e*., differences between the *R*_0_ of the two competing strains) differences in life-history traits matter away from the endemic equilibrium. In particular, faster strains have higher probabilities of fixation when the pathogen population is growing (during epidemics) and, conversely, slower strains have higher probabilities of fixation when the pathogen population is crashing. The analysis of scenarios where the epidemiological dynamics is unstable and leads to recurrent epidemics is more challenging but may lead to unexpected evolutionary dynamics (Read and Keeling, 2007).

We analyzed the effects of demographic stochasticity induced by finite population size but environmental stochasticity may also affect evolution (Frank and Slatkin, 1990; Starrfelt and Kokko, 2012; Schreiber et al., 2015). Environmental factors are known to have dramatic impacts on pathogen transmission and it would thus be particularly relevant to expand the current framework to account for the effects of random perturbations of the environment on pathogen evolution (Nguyen et al., 2015).

Another possible extension of this model would be to analyze the effect of demographic stochasticity on the multi-locus dynamics of pathogens. Indeed, the interaction between genetic drift and selection is known to yield complex evolutionary dynamics resulting in the build up of negative linkage disequilibrium between loci. But the analysis of this so-called Hill-Robertson effect is often restricted to population genetics models with fixed population size. The build up of linkage disequilibrium in some epidemiological models has been discussed in some simulation models (Althaus and Bonhoeffer, 2005; Fraser, 2005). Our model provides a theoretical framework to explore the effect of finite population size on multi-locus dynamics of pathogens and to generate more accurate predictions on the evolution of drug resistance (Day and Gandon, 2012).

Finally, although we have presented our results in the context of pathogen evolution, it is hopefully clear that a very similar theoretical framework could be used to study other examples of life history evolution in the context of demographic stochasticity. Current general life history theory largely neglects the evolutionary consequences of stochasticity arising from small population sizes. Our results suggest that it would be profitable to determine what sorts of insights might be gained for life history evolution more generally by using the type of theoretical framework developed here.

## Acknowledgements

Some of this work was done while TLP was supported by a Fondation Sciences Mathematiques de Paris postdoctoral fellowship. AL thanks the Center for Interdisciplinary Research in Biology (Collége de France) for funding. SG thanks the CNRS (PICS and PEPS MPI) for funding and Gauthier Boaglio for his help in the development of the simulation code. Simulations were performed on the Montpellier Bioinformatics Biodiversity cluster.

